# Phase separation of chromatin brush driven by enzymatic reaction dynamics of histone posttranslational modifications

**DOI:** 10.1101/2020.11.30.405134

**Authors:** Tetsuya Yamamoto, Takahiro Sakaue, Helmut Schiessel

## Abstract

The nuclei of undifferentiated cells show uniform decompacted chromatin while during development nuclei decrease in size and foci of condensed chromatin appear, reminiscent of phase separation. This study is motivated by recent experiments that suggest that the unbinding of enzymes that chemically modify (acetylate) histone tails causes decompaction of condensed chromatin. Here we take into account the enzymatic reactions of histone modifications to predict the phase separation of chromatin in a model system, the chromatin brush, which mimics chromatin at the proximity of a nuclear membrane. The model contains ‘activators’ and ‘silencers’, which change the state of the nucleosomes to (transcriptionally) active or inactive via the Michaelis-Menten kinetics. Our theory predicts that the chromatin brush will phase separate when the brush height is reduced below a threshold height. The phase separation is driven by an anti-correlation: Activators change the state of nucleosomes to the active state suppressing the binding of silencers to these nucleosomes and *vice versa*.

## 1 Introduction

In eukaryotic cells, DNA is packed into the nucleus in the form of chromatin, a complex of DNA and histone proteins^1^. The repeating unit of chromatin is the nucleosome, where DNA is wound around an octamer of histone proteins by 1.65 turns^2^. The transcription dynamics of a gene depends on the local concentration of nucleosomes^3^. In differentiated cells, heterochromatin regions, in which the local concentration of nucleosomes is relatively large, coexist with euchromatin regions, in which the local concentration of nucleosomes is relatively small. Genetic expression is suppressed in heterochromatin regions and is active in euchromatin regions. In many cases, heterochromatin regions are located near the periphery of the nucleus and euchromatin regions are located at the center of the nucleus. This canonical structure is stabilized by proteins that tether chromatin to nuclear membranes, such as lamin A/C and lamin B receptor (LBR)^4^.

In contrast to differentiated cells, the local concentration of nucleosomes in undifferentiated cells is relatively uniform^5–7^. The size of the nucleus decreases upon development and foci of heterochromatin are produced after a couple of cell divisions^6^. The transition of the chromatin structure during development is analogous to phase separation. It is therefore of interest to theoretically predict the physical mechanism involved in the symmetry breaking during the development. Theoretical efforts have been made to predict the phase separation of chromatin^8–21^. In many cases, chromatin is theoretically treated as copolymers of euchromatin- and heterochromatin-like blocks^8–16^. These models cannot therefore be used to predict the mechanism of symmetry breaking in chromatin, since the broken symmetry is already tailored into its molecular structure ^*^.

We have developed a theory of the phase separation of DNA brushes, in which DNA is end-grafted to a surface to form a polymer brush and is allowed to form a chromatin-like complex^17,18^. We treat DNA as a polymer composed of identical segments. The brush structure is a simplified representation of chromatin near the nuclear membrane. Nucleosomes are assembled at each segment at a constant rate and are disassembled when they collide with elongating RNA polymerases^23^. The brush is collapsed by the attractive interactions between nucleosomes when the nucleosome occupancy is large and swollen by the repulsive interactions between vacant DNA segments when the nucleosome occupancy is small. Our theory predicts that nucleosome disassembly as a result of transcription drives the phase separation of the chromatin brush and predicts the canonical structure of chromatin in the nucleus: Collapsed chains with large nucleosome occupany lie at the surface and swollen chains with small nuclear occupancy occupy the space above the collapsed chains. An extension of this theory predicts that the Poisson ratio of chromatin gels can be negative^19^. This prediction may correspond to the case of stem cells which feature negative Poisson ratios^24^.

The tails of histone proteins are chemically modified by enzymes, such as histone acetyltransferases (HAT) and histone methyltransferases (HMT). Chemical states of the histone tails (so-called histone marks) correlate with the dynamics of gene expression. Histone tails are acetylated (at H3K27) before transcription is initiated, implying a causal relationship between gene activation and the tail acetylation^25,26^. Nucleosomes in euchromatic regions tend to have histone marks that indicate active genes and nucleosomes in heterochromatic regions tend to have histone marks that indicate inactive genes. Proteins bind to nucleosomes depending on the chemical state of histone tails. Recent experiments have shown that HP1 proteins that bind to nucleosomes in heterochromatin show phase separation and form macromolecular condensates^27,28^. These experimental results imply that transitions of histone states play an important role in the phase separation of chromatin.

The transitions of histone states were taken into account in recent theories to predict the discontinuous transitions of chromatin and the suppression of the growth of domains due to the forced reset of histone states. These theories treat the transition of histone states by using the free energy minimization^29^ and an extension of model B^20^, and do not explicitly take into account the dynamics of enzymes that change histone states. With this approach, the transition rate does not depend on the local concentration of enzymes and on their reaction kinetics. However, recent experiments have shown that mutating MRG-1 proteins that mediate the binding of CBP/p300 (one type of HAT) to euchromatin changes the state of histones in the heterochromatin region and decompacts heterochromatin^30^. Motivated by the latter experimental result, we here take into account the binding between enzymes and chromatin and the kinetics of enzymatic reactions of histone state transitions to predict the phase separation of chromatin in a brush. For simplicity, we take into account two types of enzymes – activators and silencers. Activators provide (transcriptionally) active modifications to wild type nucleosomes and silencers provide inactive modifications to wild type nucleosomes. The phase separation of a chromatin brush is driven by an anticorrelation: Activators change the state of nucleosomes so that silencers cannot bind to them and silencers change the state of nucleosomes so that activators cannot bind to them.

## 2 Model

### 2.1 Chromatin brush

We represent chromatin at the proximity of the nuclear membrane as a polymer brush, where DNA chains are end-grafted to a planar surface with the grafting density σ and nucleosomes are assembled along these chains, see fig. 1**a**. This model is motivated by the fact that chromatin is tethered to the nuclear membrane via lamin A/C and lamin B receptor (LBR) ^4^. Each chromatin chain is composed of *N*_0_ segments of length *l_a_*. For simplicity, we assume that each segment carries only one nucleosome. Each nucleosome takes one of three states – a (transcriptionally) active state ‘Ha’, a silent state ‘Hs’, and a neutral (wild type) state ‘H’, reflecting the posttranslational modifications of its histone tails. The chromatin brush is in an aqueous solution that contains three types of enzymes – activators ‘A’, silencers ‘S’, and erasers ‘E’. Activators change nucleosomes in the neutral state to the active state and these enzymes can bind to nucleosomes in the neutral state or in the active state. Silencers ‘S’ change nucleosomes in the neutral state to the silent state and these enzymes can bind to nucleosomes in the neutral state or in the silent state. Erasers ‘E’ change nucleosomes in either the active or the inactive state back to the neutral state.

**Fig. 1.**
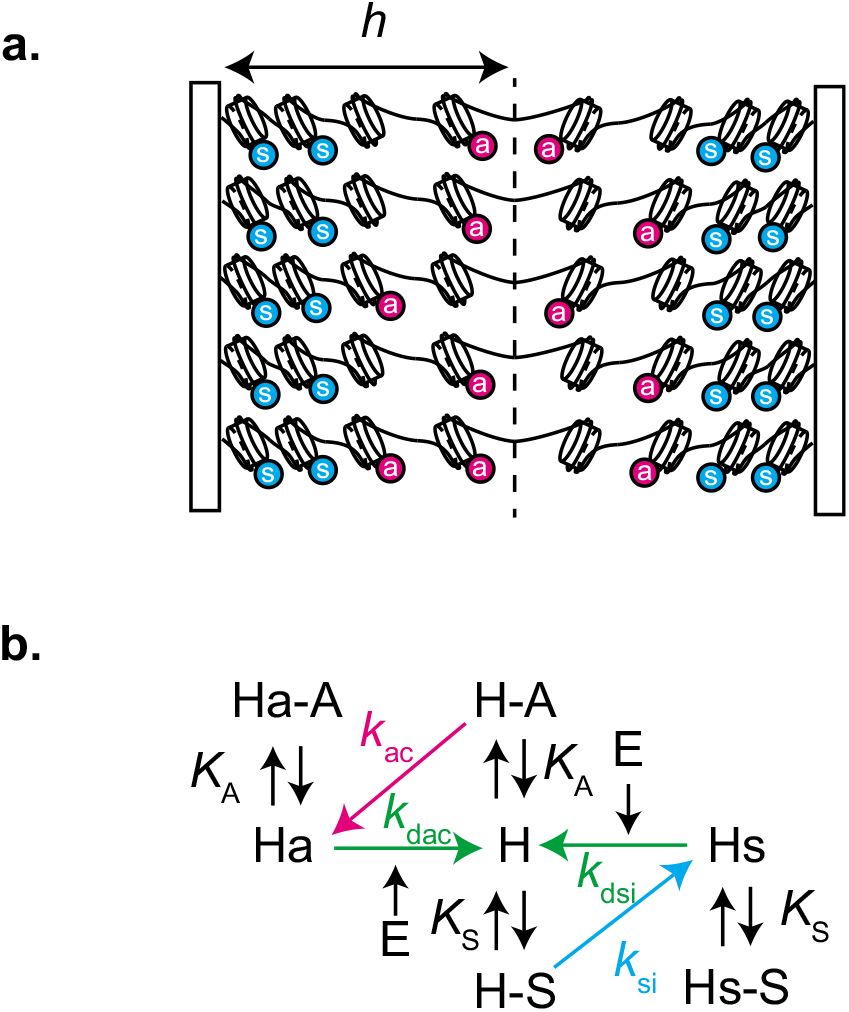
Chromatin at the proximity of the nuclear membrane is represented as a brush, where DNA chains are end-grafted to two opposing planer surfaces (**a**). We assume that each DNA segment is complexed into a nucleosome. Nucleosomes take either of three states — neutral, active, and silent, indicated by H, Ha, and Hs, respectively, in (**b**). The transitions betweeen states is driven by three types of enzymes – activators, silencers, erasers, indicated by A, S, and E, respectively, in (**b**). Activators bind to nucleosomes in either the neutral or the active state (indicated by H – A and Ha – A). Silencers bind to nucleosomes in either the neutral or the silent state (indicated by H – S and Hs – S). When phase separation occurs, the chromatin brush forms a double-layer structure where each layer corresponds to a phase.

### 2.2 Free energy

The free energy per unit area has the form

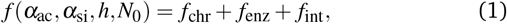

where *f*_chr_ is the free energy density of chromatin in the brush, *f*_enz_ is the free energy density due to the translational entropy of the enzymes (and solvent molecules), and *f*_int_ is the free energy due to the interactions between chromatin segments and enzymes. *h* denotes the height of the brush. α_ac_ is the fraction of active nucleosomes and *α*_si_ is the fraction of silent nucleosomes. These fractions are determined by the kinetics of the posttranslational modification of the histone tails.

For simplicity, we use the Alexander approximation, for which the local concentration of chain segment is assumed to be uniform in each phase of the brush. With this approximation, the free energy density of the chromatin brush is given by

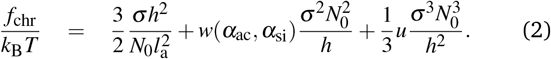

The first term of eq. (2) is the free energy density due to the conformational entropy of chromatin. The second and third terms account for the two- and three-body interactions between chain segments. The 2nd virial coefficient *w*(*α*_ac_, *α*_si_) has the form

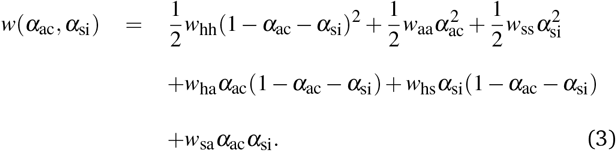

The *w*’s on the right hand side are 2nd virial coefficients. Specifically whh accounts for the (neutral nucleosome)-(neutral nucleosome) interactions, *w*_aa_ for the (active nucleosome)-(active nucleosome) interactions and *w*_ss_ for the (silent nucleosome)-(silent nucleosome) interactions. The other *w*’s account for the “nondiagonal” interactions between nucleosomes in different states. We implicitly take into account the interactions between nucleosomes via binding proteins, such as HP1, in the values (and the signs) of these 2nd virial coefficients. For simplicity, we assume that *w*_ss_ has a negative value and that *w*_0_ ≡ *w*_aa_ = *w*_hh_ = *w*_ha_ = *w*_hs_ = *w*_sa_ > 0. *u* is the 3rd virial coefficient. *k*_B_*T* is the thermal energy (*k*_B_: Boltzmann constant, *T*: absolute temperature).

The free energy density due to the translational entropy of enzymes has the form

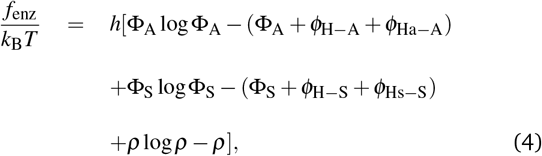

The first, third, and fifth terms are the free energy contributions due to the translational entropy of activators, silencers and erasers, respectively. The sum of the second, fourth, and sixth terms results from the free energy contributions due to the translational entropy of solvent. Φ_A_ and Φ_S_ are the local concentrations of activators and silencers that freely diffuse between chromatin in the solvent. *ρ* is the corresponding concentration of erasers. The local concentration *ϕ*_H–A_(*t*) of the neutral nucleosomes to which activators are bound and the local concentration *ϕ*_H–S_ of the neutral nucleosomes to which silencers are bound have the forms

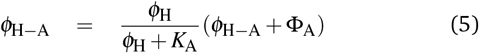

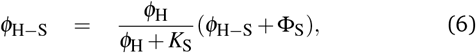

where *ϕ*_H_ (= *σN*_0_(1 – *α*_ac_ – *α*_si_)/*h*) is the local concentration of nucleosomes in the neutral state, see the black arrows in the middle column of fig. 1**b**. The occupancy of activators increases with decreasing the equilibrium constant *K*_A_. The occupancy of silencers increases with decreasing the equilibrium constant *K*_S_. Similary, the local concentration *ϕ*_Ha–A_ of the active nucleosomes to which activators are bound and the local concentration *ϕ*_Hs–S_ of the silent nucleosomes to which silencers are bound have the forms

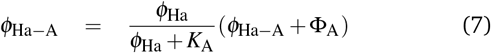

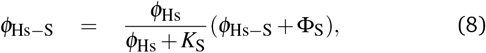

where *ϕ*_Ha_ (= *σN*_0_*α*_ac_/*h*) and *ϕ*_Hs_ (= *σN*_0_*α*_si_/*h*) are the local concentration of nucleosomes in the active and silent states, respectively, see the black arrows in the left and right columns of fig. 1**b**. For simplicity, we assume that activators bind to neutral and active nucleosomes with the same equilibrium constant *K*_A_ and that silencers bind to neutral and silent nucleosomes with the same equilibrium constant *K*_S_. Our theory implicitly takes into account proteins that mediate the binding of enzymes to chromatin in the binding constants *K*_A_ and *K*_S_.

The free energy density describing the interactions between chromatin and enzymes is given by

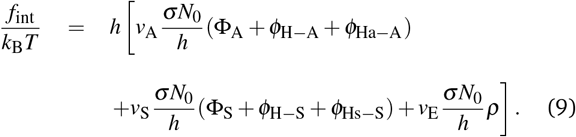

The first term of eq. (9) is the free energy density due to the interactions between activators and chromatin segments and *v*_A_ is the 2nd virial coefficient that accounts for these interactions. The second term accounts for the interactions between silencers and chromatin segments with the second virial coefficient *v*_S_. Finally, the third term describes the interactions between erasers and chromatin segments with the second virial coefficient *v*_E_. For simplicity, we set these three second virial coefficients to the same value, *v*_0_ ≡ *v*_A_ = *v*_S_ = *v*_E_. With eq. (9), we assume that the free energy due to the interactions between enzymes and chromatin segments does not depend on the nucleosomal state and on whether enzymes are bound to nucleosomes or freely diffusing. For simplicity, we neglect erasers that are bound to nucleosomes.

### 2.3 Two-phase coexistent state

In the two-phase coexistence state, the free energy per unit area has the form

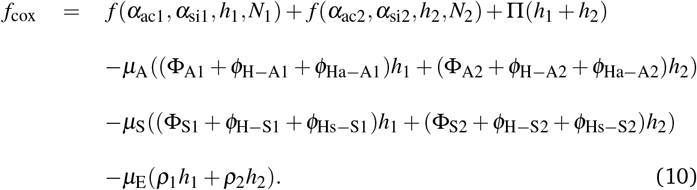

Π is the osmotic pressure in the brush. *μ*_A_, *μ*_S_, and *μ*_E_ are the chemical potentials of activators, silencers, and erasers, respectively. These chemical potentials can be used as Lagrange multipliers to fix the number of corresponding enzymes in the system.

Motivated by the fact that heterochromatin is tethered to nuclear membranes, we treat the case in which the phase-separated chromatin brush forms a double-layer, where the bottom layer is composed of one phase (with its quantities indicated by the subscript 1) and the top layer of the other phase (indicated by the subscript 2). The height of the bottom layer is *h*_1_ and it is composed of *N*_1_ segments, where the fractions of nucleosomes in the active and silent states are *α*_ac1_ and *α*_si1_, respectively. The height of the top layer is *h*_2_ and it is composed of *N*_2_ segments with *α*_ac2_ and *α*_si2_ denoting the fractions of nucleosomes in the active and silent states.

For simplicity, we assume here that the relaxation dynamics of the brush heights and the diffusion (and the binding-unbinding of enzymes to nucleosomes) are much faster than the state transitions of nucleosomes. With this assumption, the brush heights (*h*_1_ and *h*_2_) and the local concentrations of enzymes (Φ_A1_, Φ_A2_, Φ_S1_, Φ_S2_, *ρ*_1_, and *ρ*_2_) are in local equilibrium with the given fractions, *α*_ac_ and *α*_si_, of nucleosomes in the active and silent states, which are determined by slower dynamics. Note that this assumption is not essential when we analyze the steady state.

Minimizing eq. (10) with respect to the local concentrations of enzymes leads to the equality of chemical potentials between two phases

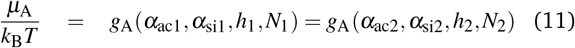

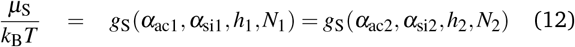

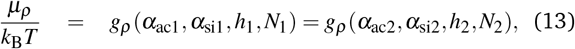

where the chemical potential functions *g*_A_, *g*_s_, and *g_ρ_* have the forms

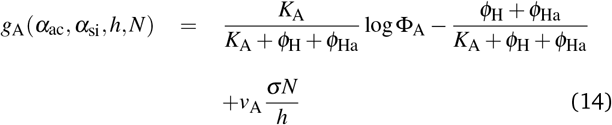

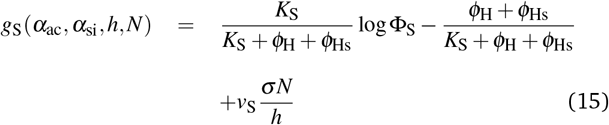

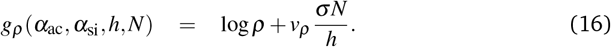

Minimizing the free energy with respect to the brush height, *h*_1_ and *h*_2_, leads to the force balance equation

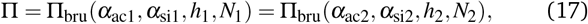

where the pressure Π_bru_ generated in the brush has the form

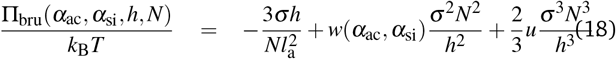

To derive eq. (18), we assumed that the local concentrations of nucleosomes is larger than the local concentrations of enzymes and thus neglected the osmotic pressure due to the translational entropy of enzymes. The height h and the local concentrations, Φ_A_, Φ_S_, and *ρ*, of enzymes in the two phases are derived for given fractions, *α*_ac_ and *α*_si_, of nucleosomes in the active and silent states by using eqs. (11) - (13) and (17).

### 2.4 Enzymatic reaction kinetics

We describe the kinetics of the enzymatic reactions of activators and silencers by using the Michaelis-Menten law. The time evolution of the state of nucleosomes has the forms

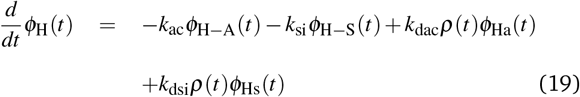

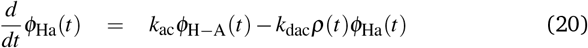

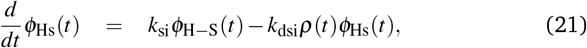

see fig. 1**b**. The first term of eq. (19) (and the first term of eq. (20)) is the transition rate of nucleosomes from the neutral state to the active state and kac is the rate constant that accounts for this process, see the magenta arrow in fig. 1**b**. The second term of eq. (19) (and the first term of eq. (21)) is the transition rate of nucleosomes from the neutral state to the silent state with rate constant k_si_, see the cyan arrow in fig. 1**b**. The third term of eq. (19) (and the second term of eq. (20)) is the transition rate of nucleosomes from the active state to the neutral state with rate constant k_dac_, see the green arrow indicated by k_dac_ in fig. 1**b**. Finally, the fourth term of eq. (19) (and the second term of eq. (21)) is the transition rate of nucleosomes from the silent state to the neutral state with rate constant *k*_dsi_, see the green arrow indicated by *k*_dsi_ in fig. 1**b**.

### 2.5 Steady state

In the steady state, *dϕ*_H_/*dt* = *dϕ*_Ha_/*dt* = *dϕ*_Hs_/*dt* = 0, eqs. (19) – (21) predict the relationships

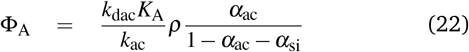

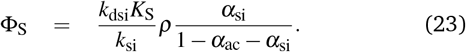

By using eqs. (11) - (13), (17), (22), and (23), we derive the height h and the fractions, α_ac_ and α_si_, of nucleosomes in the active and silent states in the steady state.

### 2.6 Characteristic scales

In the equilibrium, the height of the brush has the form

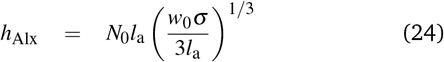

for the case in which all nucleosomes are in the active or neutral state. The concentration of nucleosomes thus scales as

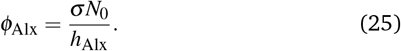

The scale of pressure is

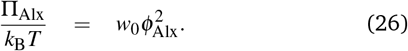

The 3rd virial coefficient *u* is rescaled by

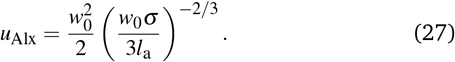

## 3 Results

### 3.1 Uniform brush with constant chemical potential

In the one phase region, the concentration of enzymes and the fractions, α_ac_ and α_si_, of nucleosomes are uniform in the brush. We first treat the grand canonical ensemble, in which the brush exchanges activators, silencers, and erasers with the exterior solution and the chemical potentials of these enzymes are constant throughout the brush (because there are no constant fluxes of these enzymes). The fractions, α_ac_ and α_si_, are derived as a function of the brush height *h* by using eqs. (11) - (13) and (22) - (23).

For cases in which the inverse equilibrium constant 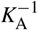 of activators is smaller than a threshold value 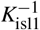 (the exterior of the region delineated by the green curves in fig. 2), our theory predicts a unique solution of nucleosome fractions, α_ac_ and α_si_, for any values of the brush height, see the cyan line in fig. 3. Whether the fractions α_ac_ and α_si_ increase or decrease with the brush height depends on the kinetic constants, the chemical potentials, and the inverse equilibrium constants of activators and silencers. When the inverse equilibrium constant 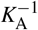 is larger than the first threshold value 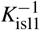 and smaller than the second threshold value 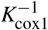 (the region between the green and the black lines in fig. 2), there is a finite window of brush heights in which two stable solutions exist, see the black line in fig. 3. However, one solution branch continues to the brush height region with a unique solution without showing an instability and it is disconnected from the other solution branch and the unstable solution. This implies that the brush height decreases without showing the transitions between the two solution branches if one starts from the brush height region with a unique solution (or the branch that does not show the instability).

**Fig. 2.**
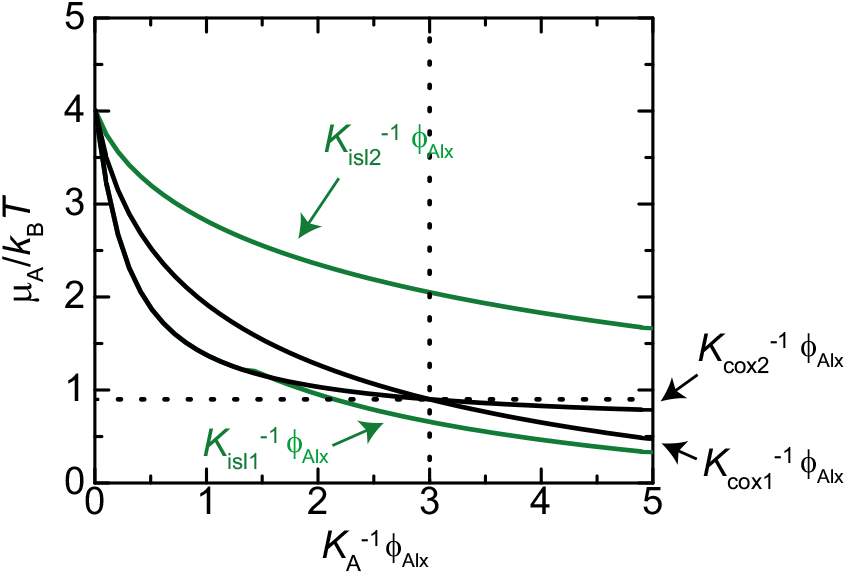
The bifurcation diagram is shown as a function of the chemical potential of activators *μ*_A_/(*k*_B_*T*) and the inverse of equilibrium constant 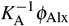. There are two stable solutions in the region delineated by the green lines and there is a transition between the two solutions in the region delineated by the black line. There is one stable solution in the exterior to these regions. The values of the other parameters used for the calculations are 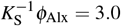, *vϕ*_Alx_ = 0.5, *μ*_S_/(*k*_B_*T*) = 0.9, *μ_ρ_*/(*k*_B_*T*) = 1.5, log(*k*_dac_*K*_A_/*k*_ac_) = log(*k*_dsi_*K*_S_/*k*_si_) = 1.5 for all lines. The values of the chemical potential *μ*_S_/(*k*_B_*T*) and the inverse equilibrium constant 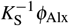 are indicated by the dotted lines.

**Fig. 3.**
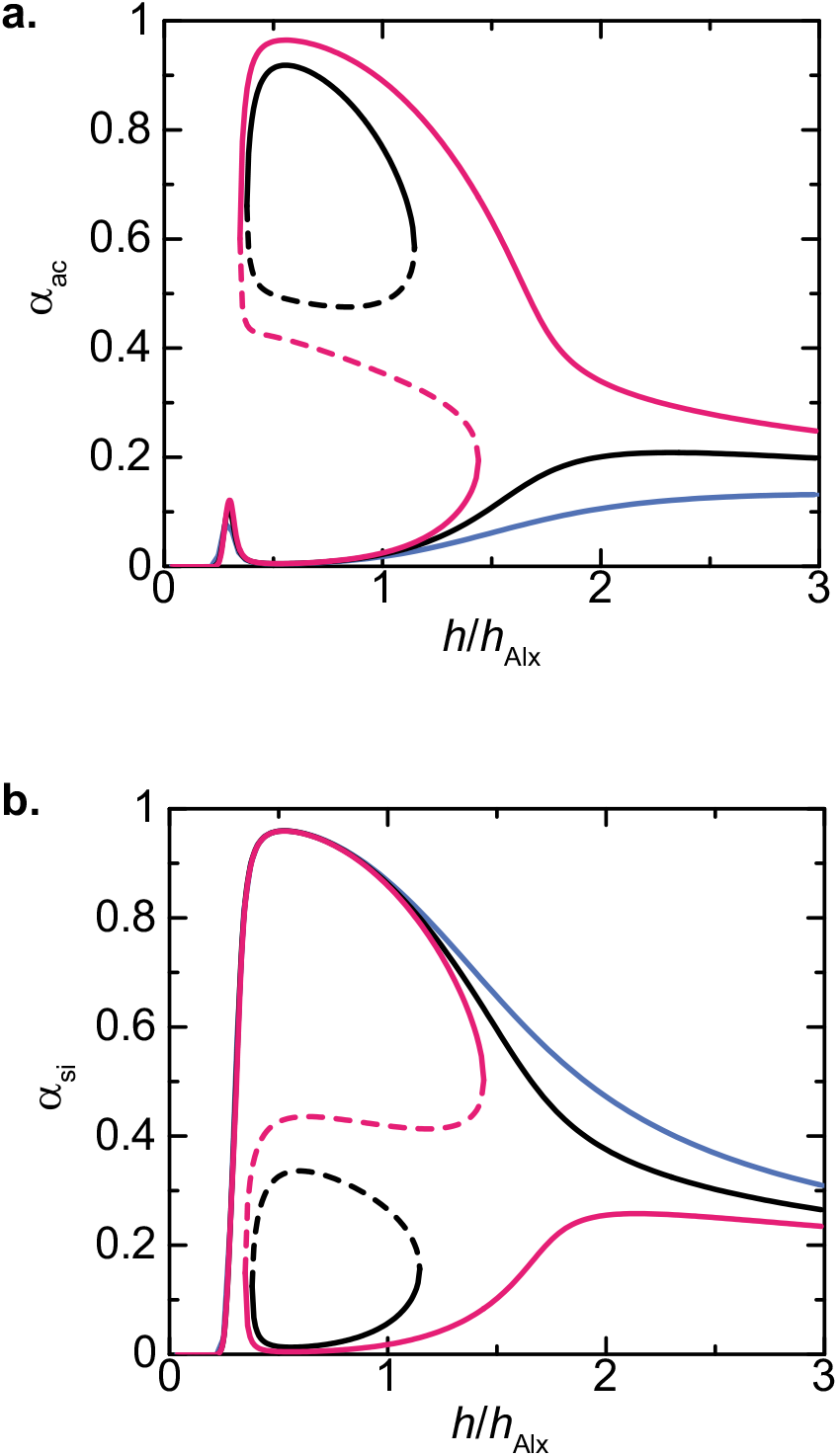
The fractions, *α*_ac_ and *α*_si_, of nucleosomes in the active and silent states are shown as functions of the brush height *h* (rescaled by *h*_Alx_) for several values of the inverse equilibrium constant 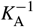 of activators, 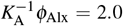 (cyan), 3.0 (black), and 3.5 (magenta). The values of the other parameters used for the calculations are 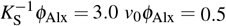, *μ*_A_/(*k*_B_*T*) = 0.8, *μ*_S_/(*k*_B_*T*) = 0.9, *μ_ρ_*/(*k*_B_*T*) = 1.5, log(*k*_dac_*K*_A_/*k*_ac_) = log(*k*_dsi_*K*_S_/*k*_si_) = 1.5 for all lines.

When the inverse equilibrium constant 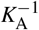 is larger than the second threshold 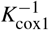 and smaller than the third threshold 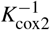 (the region delineated by the black curves in fig. 2), there is a finite window of brush heights in which two stable solutions exist, see the magenta lines in fig. 3. Each solution branch has a point at which the solution becomes unstable. The brush thus shows a transition between a euchromatin-like state in which the fraction *α*_ac_ of active nucleosomes is relatively large (and the fraction *α*_si_ of silent nucleosomes is relatively small) and a heterochromatin-like state in which the fraction *α*_ac_ of active nucleosomes is relatively small (and the fraction *α*_si_ of silent nucleosomes is relatively large). The local concentration of nucleosomes (that are the substrate of the enzymes) increases with decreasing the brush height. This increases the rate of enzymatic reactions that change the states of the nucleosomes. The bistable solutions result from an anti-correlation: Activators change nucleosomes in the neutral state to the active state so that silencers cannot bind them, whereas silencers change nucleosomes in the neutral state to the silent state so that activators cannot bind them. Because there are many parameters involved in our model, we limit our discussion to the coexistence between the two stable solutions found in this section.

### 3.2 Two-phase brush with constant chemical potential

In two-phase coexistent states, the concentrations of enzymes, the fractions, *α*_ac_ and *α*_si_, and brush heights are determined by the equality of the pressure and the chemical potentials of activators and silencers between the two phases. We here treat the phase separation of the chromatin brush in the grand canonical ensemble, in which the equality of chemical potentials is automatically satisfied because the chemical potentials are constant throughout the system (this part has been solved in sec. 3.1). We here use eqs. (17) and (18) to predict the coexistence of the two stable solutions, found in sec. 3.1.

Eqs. (17) and (18) predict that when the pressure is smaller than a threshold value Π_sp1_ or larger than the second threshold value Π_sp2_, the brush height decreases monotonically with increasing the applied pressure, see fig. 4. In a window of applied pressures, Π_sp1_ < ⊓ < Π_sp2_, there are two stable solutions. We here use the Maxwell construction

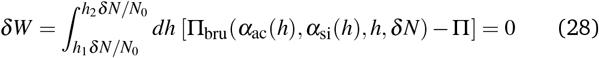

to predict the conditions at which two phases coexist. Eq. (28) states that the work *δW* necessary to change a small portion *δN* of subchains from one phase to the other is zero. The integral of eq. (28) should be performed along the unstable solutions (see the broken line in fig. 4). ‘Unstable solutions’ refers here to solutions that are unstable in either the equality of chemical potentials (eqs. (11) - (13) and (22) - (23)), the mechanical balance (eq. (17)), or both. We have previously used the same treatment to predict the phase separation of a chromatin brush^17–19^.

**Fig. 4.**
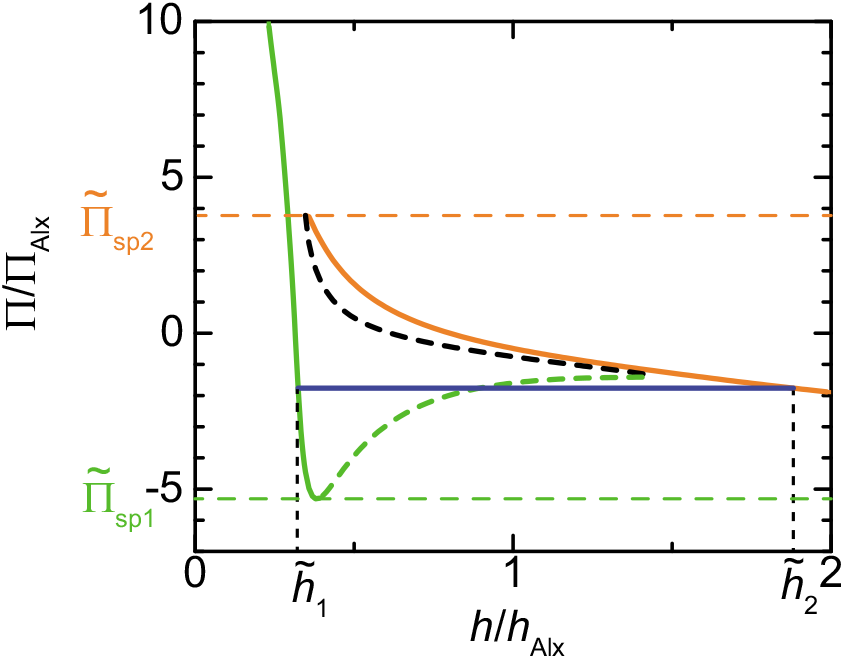
The brush height *h*/*h*_Alx_ (rescaled by the height of Alexander brush) is shown as a function of the applied pressure Π/Π_Aix_ (rescaled by the scale of pressure of Alexander brush). We used 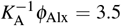, 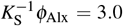, *v*_0_*ϕ*_Alx_ = 0.5, *v*_E_ϕ_Alx_ = 0.5, *μ*_A_/(*k*_B_*T*) = 0.8, *μ*_S_/(*k*_B_*T*) = 0.9, *μ_ρ_*/(*k*_B_*T*) = 1.5, log(*k*_dac_*K*_A_/*k*_ac_) = log(*k*_dsi_*K*_S_/*k*_si_) = 1.5, *w*_ss_/*w*_0_ = −2.0, and *u*/*u*_Alx_ = 0.01 for the calculations. The orange and light green solid lines correspond to the two branches of stable solutions of the magenta curve in fig. 3. The broken curves are unstable solutions.

Our theory predicts that the swollen phase, in which the height of subchains is larger, coexists with the collapsed phase, in which the height of subchains is smaller. In the swollen phase, the fraction *α*_ac_ of active nucleosomes is relatively large and the fraction *α*_si_ of silent nucleosomes is relatively small, analogous to euchromatin, see the orange curve in fig. 5. In the collapsed phase, the fraction *α*_ac_ of active nucleosomes is relatively small and the fraction *α*_si_ of silent nucleosomes is relatively large, analogous to heterochromatin, see the light green curve in fig. 5. Our theory therefore predicts the coexistence of a euchromatin-like phase and a heterochromatin-like phase. In the swollen phase, the fraction *α*_ac_ of active nucleosomes decreases and the fraction *α*_si_ of silent nucleosomes increases as the inverse equilibrium constant 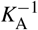 decreases. In contrast, in the collapsed phase, the fractions, *α*_ac_ and *α*_si_, do not change significantly with decreasing the inverse equilibrium constant 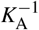. This result is not consistent with the experiments by Gasser and coworkers, which suggest that heterochromatin is decompressed by the mutation of proteins that bind activators to euchromatin^30^, at least in the two-phase coexistent state and set of parameters that are used in our numerical calculations. This may be because we here treat the grand canonical ensemble, in which the total number of enzymes is not constant. The number of activators in the swollen phase increases with increasing the inverse equilibrium constant 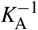. These activators are provided mainly from the external solution, rather than the collapsed phase.

**Fig. 5.**
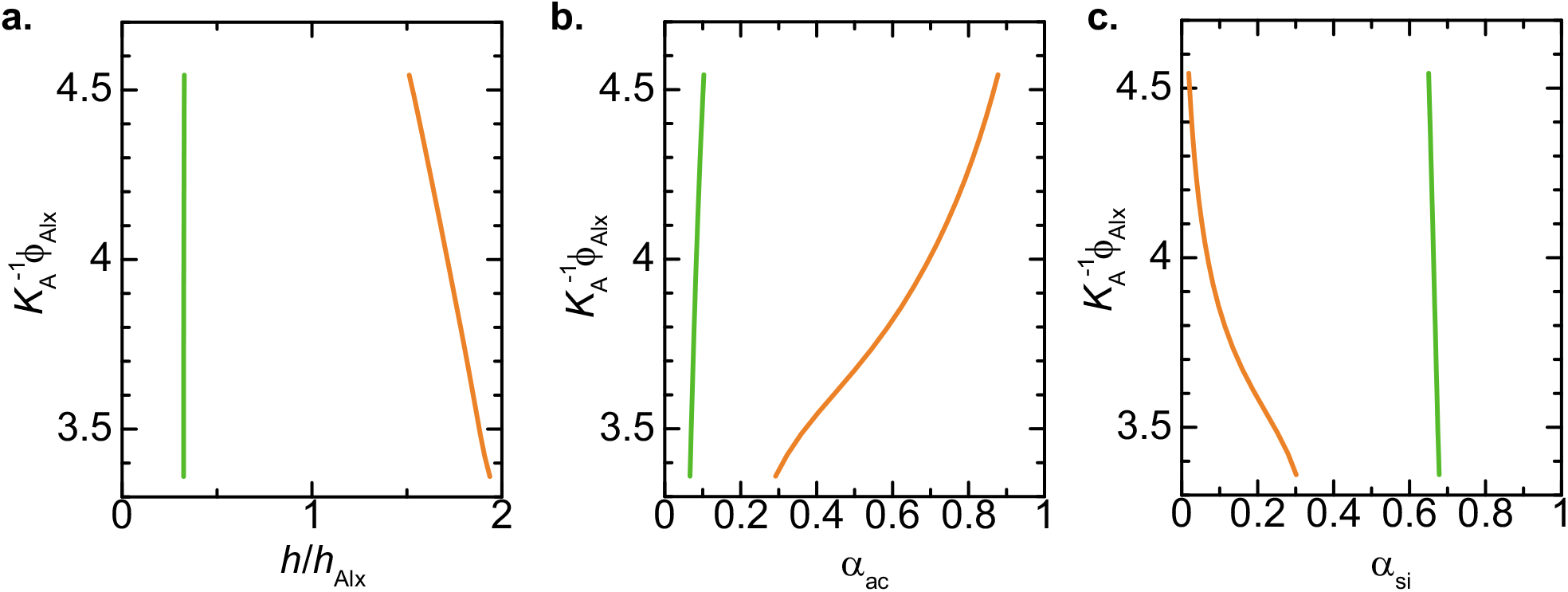
(**a**) The brush height *h* (rescaled by the height *h*_Alx_ of Alexander brush), (**b**) the fraction *α*_ac_ of active nucleosomes, and (**c**) the fraction *α*_si_ of silent nucleosomes of two coexisting phases are shown as a function of the inverse equilibrium constant 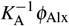. The values of collapsed phase are shown by the light green curves and the values of swollen phase are shown by the organge curves. We used 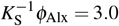, *v*_0_*ϕ*_Alx_ = 0.5, *μ*_A_/(*k*_B_*T*) = 0.8, *μ*_S_/(*k*_B_*T*) = 0.9, *μ_ρ_*/(*k*_B_*T*) = 1.5, log(*k*_dac_*K*_A_/*k*_ac_) = log(*k*_dsi_*K*_S_/*k*_si_) = 1.5, *w*_ss_/*w*_0_ = −2.0, and *u*/*u*_Alx_ = 0.01 for the calculations.

### 3.3 Two phase brush with constant number of activators and silencers

The fact that heterochromatin shows decompaction due to the release of HATs from euchromatin^30^ implies that a cell nucleus may be better modeled by the canonical ensemble, in which the number of enzymes is constant. To simplify the calculations, we here treat the case in which the numbers of activators and silencers, nA and ns, are constant in the system, but the number of erasers can change to keep the chemical potential of erasers constant throughout the system. In the two phase coexistent state, the chemical potentials of each enzyme are equal between the two phases. We derive the chemical potentials, *μ*_A_ and *μ*_S_, so that the total number of each enzyme is constant and satisfies eq. (28) by changing the brush height.

Fig. 6 treats cases in which the height of a chromatin brush of a uniform euchromatin-like state is reduced. This mimics chromatin of an undifferentiated cell, where the size of the nucleus decreases by cell division. When the inverse equilibrium constant 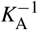 of activators is within the window 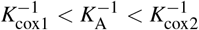, in which two solutions are stable, the chromatin brush shows phase separation for the brush heights smaller than a threshold value, see the magenta and light green curves in fig. 6. In one phase, the fraction *α*_ac_ of active nucleosomes is relatively large and the fraction *α*_si_ of silent nucleosomes is relatively small, analogous to euchromatin. In the other phase, the fraction *α*_ac_ of active nucleosomes is relatively small and the fraction *α*_si_ of silent nucleosomes is relatively large, analogous to heterochromatin. The fraction *δ* (≡ *N*_1_/*N*_0_) of heterochromatin-like phase increases as the brush height decreases. In contrast, when the inverse equilibrium constant 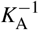 of activators is smaller than the bifurcation threshold 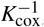, the chromatin brush does not show phase separation, see the cyan curve in Fig. 6. With the parameters that we used in Fig. 6, the fraction *α*_ac_ of active nucleosomes in the heterochromatin-like phase increases, albeit slightly, with decreasing the inverse equilibrium constant 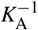, compare the magenta and light green curves. The fraction *α*_ac_ of active nucleosomes of heterochromatin increases significantly if the inverse equilibrium constant 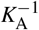 is reduced to a value smaller than the threshold 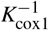, compare the magenta and cyan curves for the same height in fig. 6.

**Fig. 6.**
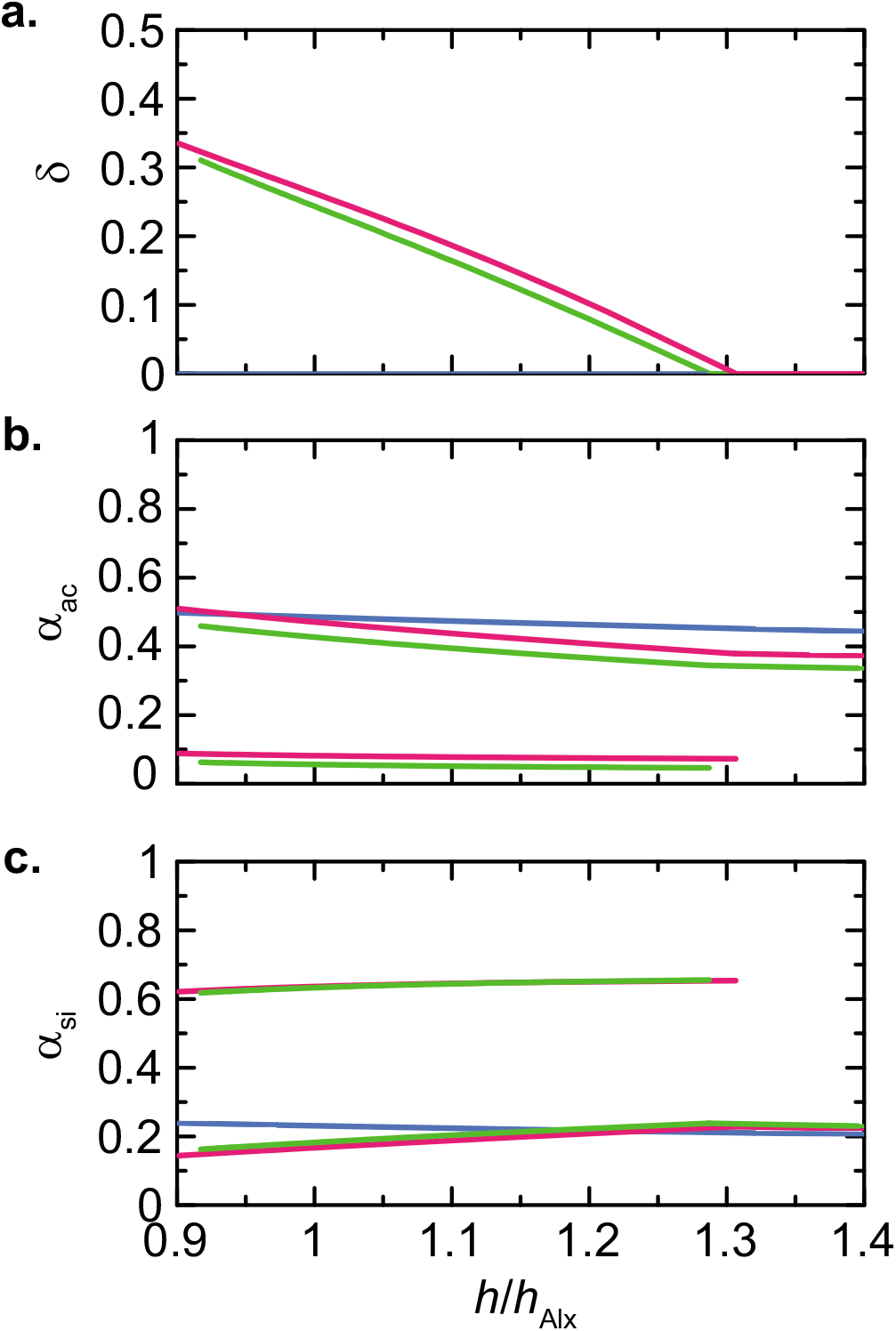
(a) The fraction *δ* of collapsed phase, (b) the fraction *α*_ac_ of active nucleosomes, (c) the fraction *α*_si_ of silent nucleosomes are shown as a function of the total brush height *h*/*h*_Alx_ (*h* = *h*_1_ + *h*_2_) for 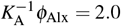 (cyan), 3:5 (magenta), and 4:5 (light green). The numbers of activators and silencers, *n*_A_ and *n*_S_, per unit area are fixed to log(*n*_A_/*h*_Alx_) = 3.97 and log(*n*_S_/*h*_Alx_) = 3.23, respectively. We used 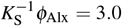, *v*_0_*ϕ*_Alx_ = 0.5, *μ_ρ_*/(*k*_B_*T*) = 1.5, log(*k*_dac_*K*_A_/*k*_ac_) = log(*k*_dsi_*K*_S_/*k*_si_) = 1.5, *w*_ss_/*w*_0_ = −2.0, and *u*/*u*_Alx_ = 0.01 for the calculations.

## 4 Discussion

Our theory treats the phase separation of a chromatin brush caused by the enzymatic reactions of histone posttranslational modifications. This theory is motivated by the fact that heterochromatin is decompacted when HAT (an activator) is unbound from euchromatin^30^. The feature of our model is that the binding of enzymes to chromatin is explicitly taken into account. Our theory predicts that the phase separation of chromatin is driven by an anti-correlation: Activators change the state of nucleosomes to the active state so that silencers cannot bind. On the other hand, silencers change the state of nucleosomes to the silent state so that activators cannot bind. This may be checked by eliminating the anti-correlation, namely by allowing activators and silencers to bind to nucleosomes in any of the states. In the latter case, eqs. (11) and (12) predict that the local concentrations of enzymes, Φ_A_ and Φ_S_, do not depend on *α*_ac_ and *α*_si_ of active and silent nucleosomes (all *ϕ*_H_ + *ϕ*_Ha_’s in eq. (14) and all *ϕ*_H_ + Φ_Hs_’s in eq. (15) should be replaced by *ϕ*_H_ + *ϕ*_Ha_ + *ϕ*_Hs_). Eqs. (22) and (23) thus have only one solution

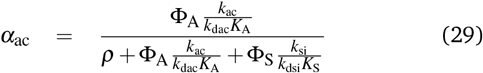

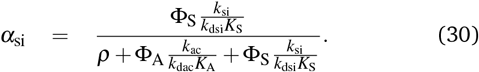

Indeed, the anti-correlation only causes bistability of nucleosome states: In one solution, the fraction *α*_ac_ of active nucleosomes is relatively large, and the fraction *α*_si_ of silent nucleosomes is relatively small, analogous to euchromatin. In the other solution, the fraction *α*_ac_ of active nucleosomes is relatively small and the fraction *α*_si_ of silent nucleosomes is relatively large, analogous to heterochromatin. The two solutions are spatially separated due to the attractive interactions between silent nucleosomes (via binding proteins, such as HP1^27,28^), see eq. (3).

Our theory predicts that the phase separation of the chromatin brush is driven by decreasing the distance between the grafting surfaces, see fig. 1). It is consistent with the fact that the phase separation of chromatin in an undifferentiated cell is driven after a couple of cell division, by which the volume of the nucleus decreases. The fraction of enzymes that bind to chromatin increases with decreasing the brush height (equivalently, increasing the nucleosome concentration), see eq. (5) - (8). This enhances the anti-correlation, discussed in the last paragraph, and stabilizes the coexistent state of euchromatin-like and heterochromatin-like phases. This prediction may be tested experimentally by detecting the time at which heterochromatin foci form on undifferentiated cells in which the proteins that bind HAT to chromatin are perturbed.

The mechanism of phase separation, demonstrated by our theory, is very different from our previous theories^17,18^ and other theories^8–16,20^. Our previous theories^17,18^ predict that the applied pressure stabilizes a condensed chromatin phase due to the attractive interactions between nucleosomes and that RNA polymerase (which destabilizes the condensed phase) is excluded from this phase due to the excluded volume interactions between RNA polymerase and nucleosomes. Some other theories also emphasize the roles played by chromatin compaction on the phase separation of chromatin^29^. Indeed, our model takes into account the attractive interactions between silent nucleosomes, see eq. (3), and the excluded volume interactions between enzymes and nucleosomes. However, with our present model, the excluded volume interactions between enzymes and nucleosomes do not play essential roles in the coexistence between the euchromatin-like phase and the heterochromatin-like phase, as defined by the two solutions of eqs. (11) - (13) and (22) - (23).

Experiments by Gasser and coworkers ^30^ and our theory imply that transcription is regulated by the localization of HAT or other enzymes to posttranslationally modify the chemical states of histone tails. This concept may be better demonstrated by simpler models in which the number of involved parameters is smaller than our present model (even if it may be less realistic than our present theory). One of the complexities of our present model is that this theory takes into account the asymmetry of the extent of compaction, the kinetic constant of histone modification, and the binding affinity (represented by the equilibrium constant in our theory) of enzymes to nucleosomes between the euchromatin-like solution and the heterochromatin-like solution. It is tempting to assume that the reaction kinetics of activators and silencers is symmetric. However, in such cases, the window of the inverse equilibrium constant, 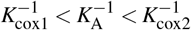, in which euchromatin-like and heterochromatin-like solutions are stabilized, converges to a point, see fig. 2. In reality, there is one more asymmetry: CpG islands of some of the genes are methylated and those of the other genes are not methylated ^22^. The binding affinity of enzymes to nucleosomes, the kinetic constant of histone modification, and the second virial coefficient may depend on the methylation state of CpG islands. Elucidating the physics of the symmetry breaking of the chromatin structure is important for understanding the transcription regulation at the early stage of development.

## Acknowledgements

This work was supported by JSPS KAKENHI Grant Number 18K03558 (T.Y.), by MEXT KAKENHI Grant Number 19H05259 (T.Y.) and JP18H05529 (T.S.), and the Deutsche Forschungsgemeinschaft (DFG, German Research Foundation) under Germany’s Excellence Strategy – EXC 2068 – 390729961–Cluster of Excellence Physics of Life of TU Dresden (H.S.). T.Y. is grateful to the fruitful discussion with Hiroshi Kimura (Tokyo Institute of Technology) and Akatsuki Kimura (National Institute of Genetics). This research was supported in part by the National Science Foundation under Grant No. NSF PHY-1748958 and NIH Grant No. R25GM067110.

## Footnotes

† Electronic Supplementary Information (ESI) available: [details of any supplementary information available should be included here]. See DOI: 10.1039/cXsm00000x/

* Indeed, CpG islands of a fraction of DNA are methylated in an early stage of development and the symmetry is already broken at this point^22^. However, for simplicity, we neglect it in the first step, assuming that DNA methylation does not play significant role in driving the phase separation of chromatin.

